# Oculomotor routines for perceptual judgments

**DOI:** 10.1101/2023.09.27.559695

**Authors:** Avi M. Aizenman, Karl R. Gegenfurtner, Alexander Goettker

**Affiliations:** Psychology Department, Giessen University, Giessen, Germany

**Keywords:** virtual reality, eye tracking, oculomotor routines

## Abstract

In everyday life we frequently make simple visual judgments about object properties, e.g., how big or wide is a certain object? Our goal is to test whether there are also task specific oculomotor routines which support perceptual judgments, similar to the well-established exploratory routines for haptic perception. In a first study, observers saw different scenes with two objects presented in a photorealistic virtual reality environment. Observers were asked to judge which of two objects was taller or wider while gaze was tracked. All tasks were performed with the same set of virtual objects in the same scenes, so that we can compare spatial characteristics of exploratory gaze behavior to quantify oculomotor routines for each task. Width judgments showed fixations around the center of the objects with larger horizontal spread. In contrast, for height judgments, gaze was shifted towards the top of the objects with larger vertical spread. These results suggest specific strategies in gaze behavior that presumably are used for perceptual judgments. To test the causal link between oculomotor behavior and perception, in a second study, observers either could freely gaze at the object or we introduced a gaze contingent set up forcing observers to fixate specific positions on the object. Discrimination performance was similar between free gaze and the gaze contingent conditions for width and height judgments. These results suggest that although gaze is adapted for different tasks, performance seems to be based on a perceptual strategy, independent of potential cues that can be provided by the oculomotor system.

## Introduction

In natural behavior, our visual system samples information actively from the environment by frequently changing gaze position. Eye movements allow the alignment of the small, highest visual acuity foveal region on objects and locations of interest. A number of factors contribute to the selection of gaze targets (see the review from Schütz, Braun, & Gegenfurtner, 2011). Some previous work has considered the ways which gaze is driven by the saliency of the stimulus. Psychophysical evidence has shown bottom-up differences in image properties between locations that were fixated or randomly chosen (Tatler, Baddeley, & Gilchrist, 2005). Subsequent work revealed higher luminance contrast or edge density at fixated regions (Parkhurst & Niebur, 2003; Baddeley & Tatler, 2006). Models have been proposed to predict such bottom-up, stimulus-driven features, to predict where observers are likely to fixate (Itti, Koch, & Niebur, 1998; Koch & Ullman, 1985). These saliency map representations act as a collection of different feature maps, each of which represent a single visual feature (such as color or size). These maps are then combined into a single map, representing saliency over the scene (Itti & Koch, 2000). Extensions of this work have created meaning maps, which represent the semantic complexity of a scene to predict gaze (Henderson & Hayes, 2017), or relied on image processing by deep neural networks to predict scan paths (Kümmerer, Bethge, & Wallis, 2022).

In contrast to this bottom-up approach, the allocation of gaze is also influenced by the given cognitive task (Yarbus, 1967), visuomotor task (Land, Mennie, & Rusted, 1999; Hayhoe, 2000; Land & Hayhoe, 2001; Rothkopf, Ballard, & Hayhoe, 2007; Sullivan, Ludwig, Damen, Mayol-Cuevas, & Gilchrist, 2021; Brouwer, Franz, & Gegenfurtner, 2009; Schütz, Braun, & Gegenfurtner, 2011), and perceptual tasks (Toscani, Valsecchi, & Gegenfurtner, 2013a). Pioneering work by Yarbus (1967) showed specific changes in gaze behavior with cognitive task. Observers were asked to perform different cognitive tasks with the same image. Not only did the objects fixated differ depending on the given task, but the locations fixated within given objects showed task-specific differences suggesting that an observer’s unique pattern of gaze carries a signature that can identify the task they are performing. The influence of task on gaze can even be seen at the scale of fine eye movements: Recent work has shown that ocular drifts during fixations aren’t random, but systematically vary by task to improve performance (Benedetto & Kagan, 2023; Lin, Intoy, Clark, Rucci, & Victor, 2023). The importance of task on gaze control has also been shown in the visuomotor domain. Rothkopf, Ballard, & Hayhoe (2007) showed that when observers were instructed to avoid objects while walking along a virtual path, participants looked towards the edge of virtual objects. When the task was to collect the object, gaze was biased towards the object’s center. These patterns were remarkably consistent across observers, and the task could even be predicted based on the fixation data. Additionally, observers fixate on different locations when asked to grasp an object rather than look at it, behavior which reflects task-specific fixation biases (Brouwer et al., 2009).

The influence of the task becomes especially relevant when looking at fixation behavior in the natural world. This early and influential work used mobile eye trackers to record gaze during everyday tasks (Land et al., 1999; Land & Hayhoe, 2001; Hayhoe & Ballard, 2005). In these experiments, gaze was tracked while observers made a sandwich or prepared tea. During such tasks, gaze is not driven by saliency, but seems to fulfill different functions, which include locating (determining the positions of objects for future use), directing (establishing target direction before initiating contact), guiding (overseeing the relative movements of multiple objects), and checking (confirming whether a specific condition is met, before completing an action). This means that the timing and decision of where we choose to look are intimately linked with behavior and action.

The evidence across tasks and modalities suggests fixations are purposeful, and that where one fixates can affect perception and decision-making. The question of how people fixate and how these locations relate to the task being performed is critical to understand how information is sampled from the visual environment, ultimately guiding visual decision-making. However, a basic taxonomy for how fixations are distributed for fundamental visual tasks is missing. In the haptic domain, Lederman et al., (1967) successfully pioneered an approach to quantify motor routines for haptics perception. They found that behavior is adapted for different tasks and that observers execute stereotypic routines when making haptic judgments. When judging an object’s texture, observers will tend to stroke the object, while when judging the hardness of the same object, observers will prod the object. When these stereotypic hand movements are constrained, performance on a matching task is reduced (Lederman & Klatzky, 1987). This means there are task-specific motor routines for haptic judgments and that these routines are optimized and even necessary for haptic exploration.

For visual perception Toscani et al., (2013a,b) developed an approach to measure the effect of fixations on lightness perception. Observers were asked to adjust the lightness of a small patch on a monitor to match a uniformly colored object. Despite variations in luminance over the object’s surface, caused mainly by shading, observers were able to match the reflectance of the objects’ surface. Gaze patterns revealed that observers didn’t fixate on the object randomly but instead fixated on brighter regions, which corresponded to their lightness match. Interestingly, observers rarely fixated on the darkest regions of the object. In a follow-up study, observers were forced to fixate on a dark or light region of the object in a gaze-contingent setup. If observers failed to fixate on the assigned location the object would disappear. When forced to fixate a darker region of the object, the match was adjusted to be darker than when observers fixated brighter regions on the object. This finding highlights a causal link between the luminance of the regions fixated, and the lightness judgment of the whole object. This work showed that fixating on the brightest region is a computationally optimal strategy for estimating lightness.

Inspired by this previous work, we want to measure whether task-specific oculomotor routines exist for common visual judgments about object properties. These judgments include surface shape (e.g. width and height) as well as surface properties (e.g. brightness). There is a wealth of evidence that humans are excellent at estimating spatial distance, which is crucial for planning more complex action sequences (such as estimating whether one’s car will fit into an empty parking spot). Prior work has shown that humans are adept at estimating length correctly and that these judgments are remarkably stable over time (Stevens & Galanter, 1957; Verrillo, 1983). Furthermore, confidence judgments track performance in spatial discrimination tasks (Baranski & Petrusic, 1994; Duyan & Balci, 2020). Previous work has additionally shown that when judging the size of an object with both visual and haptic input, vision often dominates the ultimate percept (Rock & Victor, 1964; Hay, Pick Jr, & Ikeda, 1965). Subsequent work showed that visual dominance occurs when the variance associated with the visual estimation is more reliable than the haptic input (Ernst & Banks, 2002). These findings emphasize the fine-tuned accuracy of visual judgments for basic object properties. Nevertheless, the precise contributions of fixations to this decision-making process remain unclear.

The measurements from previous studies that quantified length and spatial distance estimations were obtained in controlled laboratory settings while observers viewed static images, with the head restrained in a chin rest, and fixation stable on a target. This is in contrast to the natural environment, where the head and eyes move freely, and we engage with the natural world for longer periods. The human visual system is active by nature (Ballard, 2021; Findlay & Gilchrist, 2003; Noë & O’Regan, 2001; Gegenfurtner, 2016), and virtual reality (VR) provides a well-controlled experimental setup while giving the observer freedom of movement in a naturalistic 3D setting. This improves the sense of presence, even in a virtual /non-physical environment. (Mestre, Fuchs, Berthoz, & Vercher, 2006). Advances in VR technology and eye tracking have made it possible to track gaze in 3D space and to map fixations to the virtual world (Clay, König, & Koenig, 2019). Although task-specific fixation biases have been documented in controlled laboratory settings, it is unclear how finely tuned fixations are to support basic visual judgments for natural object properties. The combination of VR and eye tracking could inform how the visual system selects relevant visual information from a naturalistic stimulus to support visual judgments. Our goal is to investigate the strategic fixation pattern executed for a given perceptual task in a naturalistic environment. Ultimately, these measurements could inform a systematic taxonomy of the functions gaze serves during visual judgments. We hypothesize that observers will use different patterns of fixation depending on the given task. One hypothesis is that observers use gaze as a measuring tape to estimate surface size properties based on additional cues related to the executed eye movement: If asked to judge an object’s height, observers may look toward the top and bottom of the object to determine the absolute height of the object. Alternatively, for surface properties such as lightness, we expect observers to look at task-relevant object locations (such as the brightest regions of the object) to inform visual judgments, a finding which would replicate previous work (Toscani et al., 2013a; Toscani, Valsecchi, & Gegenfurtner, 2013b). In a first experiment, observers were shown two objects in photorealistic VR and were asked to make visual judgments about these objects across a range of object properties. In a second experiment, a gaze-contingent paradigm was used to limit fixation to specific locations on the object. By interrupting the preferred fixation strategy, we can measure if there is a causal link between oculomotor behavior and perceptual judgments.

## Experiment 1: Two Objects

## Methods

### Participants

Fifteen people participated in the experiment. They were 21-31 years of age (twelve identified as female, three identified as male). The experimental protocol was approved by the Institutional Review Board at our university (LEK-2021-0028) in accordance with the Declaration of Helsinki. Participants signed informed consent forms before participating.

### Apparatus

The experiment was presented using the HTC Vive Pro Eye headset, which includes a built-in eye tracker (Tobii XR). The Tobii XR SDK (Tobii, 2020) and Vive SRanipal SDK (Vive, 2020b) were used to access eye tracking data at 90Hz. According to the manufacturer, tracking accuracy is *∼* 0.5 – 1.1° (Vive, 2020a). The HMD includes two OLED screens, with a resolution of 1400*×*1600 pixels per eye. The experiment was run on a PC with Windows 10 64-bit operating system, AMD Ryzen Threadripper 3990X 64-core processor with 2.9GHz, 256GB RAM, and an NVIDIA GeForce RTX 3090 graphics card. The experiment was developed using Unity (version 2019.3.8f1), and the experiment frame rate reached ~90Hz.

### Stimuli

Objects were selected from the ArchVizPRO (interior Volume 6) asset pack, available on the Unity Asset Store. These objects are specifically designed to closely resemble real-world materials and textures with a high degree of realism. Sixteen objects from this set were selected for the experiment, as well as a table for the objects to sit on.

### Experiment Design

At the beginning of the experimental session, participants placed and adjusted the HMD on their head to a comfortable position that enabled a full field of view. They also adjusted the separation between the left and right screens to match the inter-ocular distance. The eye tracker was then calibrated using the five-point calibration procedure provided by the Vive Pro Eye. To confirm that the calibration was successful and eye tracking error minimized, we developed our own calibration check. A small target was displayed at different positions in the central visual field, and participants were instructed to fixate the target center and press the spacebar once they thought fixation was accurate. The targets were displayed at virtual distances of 1.5m. The targets were shown in random order in nine positions (in a 3 by 3 grid); the central position was presented at eye-level straight ahead. The targets were separated 10° horizontally and vertically from each other. If the error between the participants’ gaze and the cube’s center was less than 1°, the experiment began.

In the experiment, two objects appeared on top of a table in a virtual reality environment (see Figure 1). At the start of each trial, observers were instructed to judge which object was taller, wider, or brighter. To respond, observers pointed a controller at the object to be selected and pressed the trigger button on the controller to choose that object. Each scene contained two objects, and in half the trials, one of the objects was shown on top of another object, such as a box (‘boosted’ trials). These boosted trials contained the same objects as the ‘un-boosted’ trials. When judging height, observers were instructed to judge the height of the object, ignoring the contribution of the booster. We introduced the boosted trials to force observers to engage in more complex oculomotor behavior, even though the objects to be judged are identical in trials with both un-boosted and boosted objects. Observers were presented with 48 trials in total, 16 trials for each task (height, width, and brightness judgments), and half of the trials contained ‘boosted’ stimuli. Tasks were performed with the same set of virtual objects in the same scene, which allows a direct comparison of the gaze characteristics for each task.

**Figure 1:**
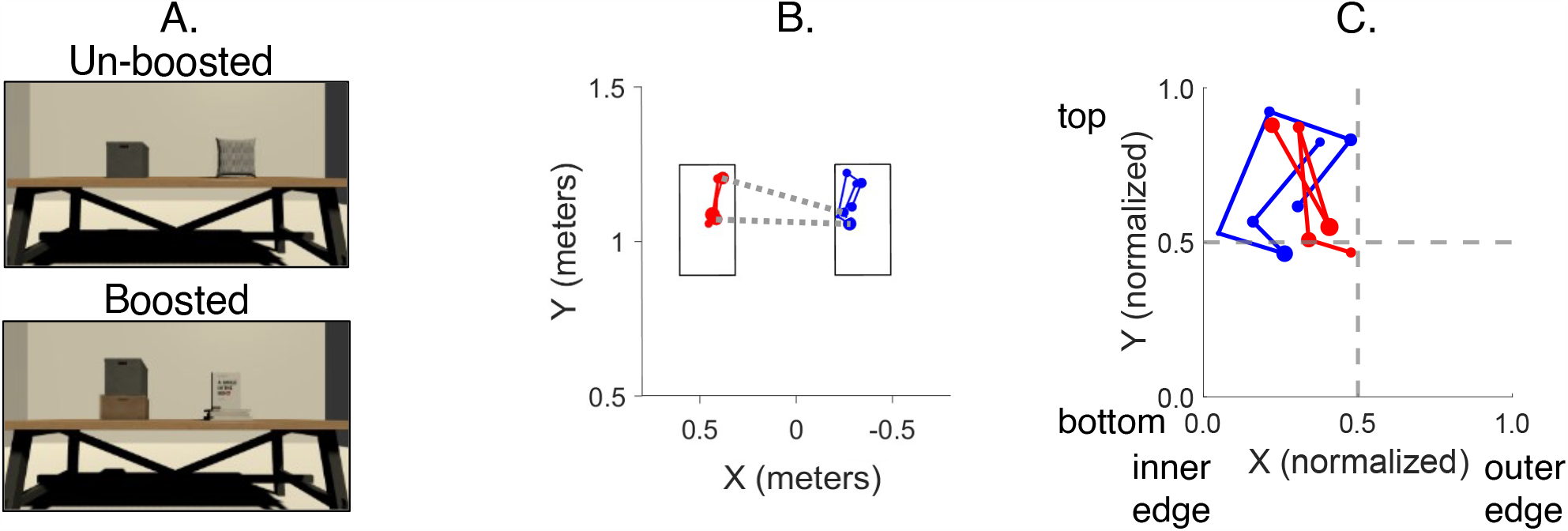
Stimuli and task in Experiment 1. A) Shows examples of trials featuring un-boosted (top panel) and boosted (bottom panel) stimuli. B) shows fixations over the two objects from a single trial. Fixations on the object are shown in virtual meters. Red data points correspond to fixations on the left object in the scene, blue points to fixations on the right object in the scene. Data markers represent fixations and the dotted grey lines between the objects show switches (saccades made between the objects). C) An example of the data shown in panel B) following the normalization procedure.

### Data Analysis

Eye tracking in virtual reality, and defining gaze vectors relative to objects provides a unique set of challenges (Clay et al., 2019). When designing the virtual environment, each object was covered by an invisible ‘collider’ with a unique label. The collider is an invisible box that defines a region of interest for a single object. The gaze data from the HTC Vive Pro Eye returns raw data in XYZ coordinates that correspond to the gaze hit point on the object (in meters) in the virtual world. Each eye-tracking sample is accompanied by the label of the collider/object that the gaze vector intersected with. At the start of every trial, the location and size of the colliders for all objects in the scene were saved in world coordinates. By saving gaze position in the same coordinates, we can map where the gaze vector (in world coordinates) intersects on the object.

Raw gaze data in XYZ coordinates, the collider label, and object location details were saved during the experiment and were analyzed offline using Matlab (Mathworks, Natick M.A.). The gaze position data was filtered with a Loess smoothing approach.

Afterward, eye velocity was computed over consecutive samples. Saccades were detected as velocity *>* 35 °/sec. To estimate how fixations are distributed along the object, we implemented a normalization procedure for gaze data samples that were part of a fixation and landed on either of the two objects in the current trial. Fixations on the edge of objects, where samples fell both on the object, and on the surround were only counted if 75% of the samples from the fixation landed on the object. A fixation was only counted if the duration was greater than 100 msec. Fixations on each object were normalized between 0-1 based on the location and size of the current object. Therefore, if a gaze sample fell exactly on an object’s center, the XY gaze sample would be normalized to (0.5,0.5) regardless of the size of the fixated object. As Figure 1B shows, fixations consistently tended to be biased towards the inner edges closest between the two objects. When averaging fixations over the two objects to identify gaze strategies, the horizontal component of gaze would obscure this edge bias. To avoid this, the left object’s X gaze position was inversed, as shown in Figure 1C. Trials were excluded from the analysis based on whether the saccade algorithm detected a saccade larger than 100 msec. This indicates a failure of the detection algorithm and limits our ability to correctly detect which gaze samples are part of a fixation. Out of 765 trials, 639 (83%) were included for analysis.

In order to visualize the distribution of gaze, heat maps were generated from the normalized gaze data. The heatmaps show fixation relative to the proportion of the object. To generate the heatmaps, normalized gaze coordinates between 0 and 1 were smoothed and aggregated into 0.025 × 0.025 bins. The varying colors show different levels of gaze density across the object. Red regions indicate regions where the participant spent time fixating while cooler colors represent regions where observers spent less time viewing the object.

For the brightness task, observers were instructed to judge which of the two objects was brighter. To analyze the average brightness of fixated pixels, we began by calculating the brightness intensity of the objects. We converted screenshots of the original objects used in the experiment to grayscale luminance intensity images using the rgb2gray Matlab function. The output is a 2D array that represents the intensity of the image at each pixel location. These images were filtered with a 21×21 pixel Gaussian filter, providing an estimate of brightness (represented as RGB intensity) over the object. For each fixation on the object, we utilized these intensity images to determine the brightness of the pixel fixated.

To assess the distribution of gaze over the object, in the horizontal and vertical direction, we analyzed the X and Y component of gaze. To quantify the behavior, we computed the median as well as the spread (difference between the 2.5 and 97.5% of the X or Y distribution). In addition, to quantify typical saccade and fixation behavior, we computed the average fixation duration, saccade amplitude, and the switch rate between objects. For all metrics, we separately computed the average across trials for each of the three conditions (width, height, and brightness judgments) for each participant. We looked for systematic differences in the data by computing repeated-measures ANOVAs with two factors: condition (width, height, or brightness) and boosted status (boosted vs un-boosted). If there was a main effect, post-hoc t-tests evaluated differences between conditions, and in order to avoid the multiple comparison problem, we report the adjusted p-value using a Holm correction. All statistical analyses were performed in JASP (JASP Team, 2023).

## Results Experiment 1

The goal of this experiment was to investigate natural gaze behavior in photorealistic VR to quantify differences in fixation characteristics for height, width, and brightness judgments. As all tasks were performed with the same set of virtual objects in the same scenes, we can compare spatial characteristics of exploratory gaze behavior to quantify oculomotor routines for each task.

To visualize overall gaze patterns, figure 2 shows average heat map representations, depicting the distribution of gaze across the objects for width, height, and brightness judgments. In these plots, values towards the top left region indicate a gaze bias towards the top of the object and towards the center edge closest to the other object. Larger values towards the bottom right region indicate fixations directed toward the bottom outer edge of the object. From these visualizations, we can see that observers did not follow a measuring-tape approach. For that, we would have expected distinct clusters at the horizontal or vertical edges of the object. Instead, we observe gaze clustering at different parts of the object. For height judgments, gaze is biased toward the top of the object. Conversely, when making width and brightness judgments, fixations tend to be biased towards the center of the objects. Interestingly, across tasks, the X component of gaze shows a bias towards smaller values, suggesting observers tend to fixate towards the inner edge of the objects to compare the parts of the object that are closest to one another.

**Figure 2:**
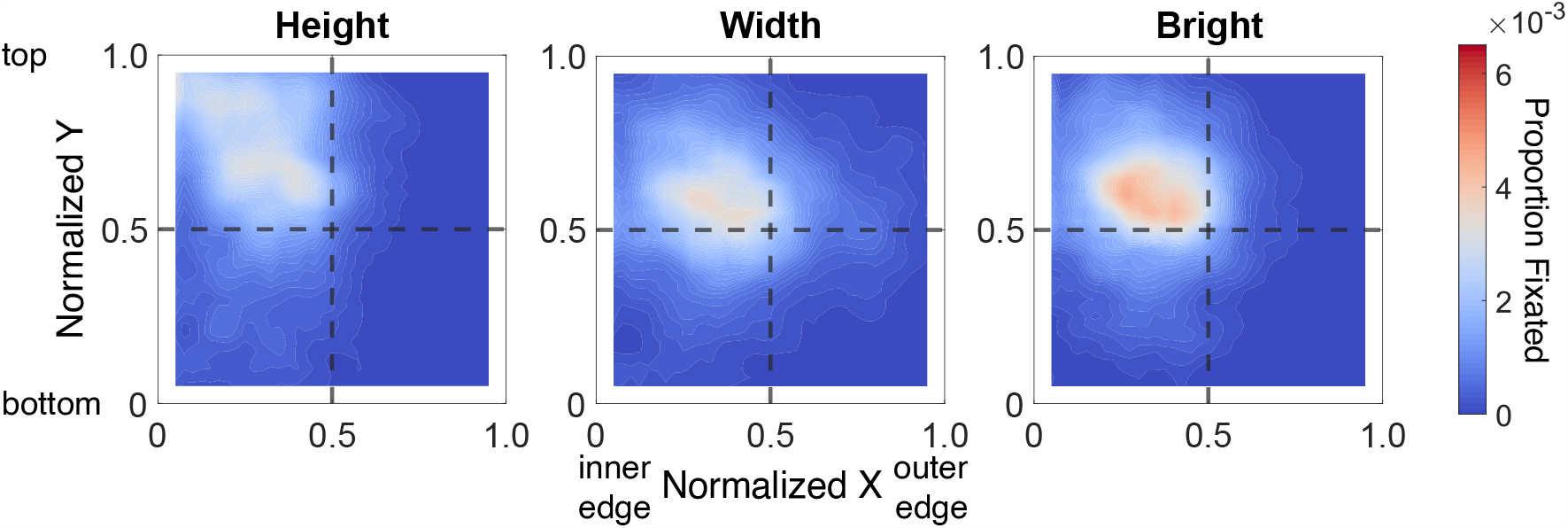
Heatmap visualizations of gaze during width, height, and brightness judgments. The heatmaps shown represent gaze relative to the object proportion. The horizontal axis shows normalized X gaze values. Smaller values represent gaze toward the center edge of the objects (the sides of the objects closest to each other), and larger values show the outer edge of the objects. The vertical axis represents normalized Y gaze values; smaller values correspond to gaze toward the bottom of the object while larger values the top of the object. The dotted grey lines represent X = 0.5 and Y = 0.5, the halfway point. The intersection of the dotted grey lines shows the object center; data that fall along this intersection represent fixations biased towards the object center. Areas with more gaze samples are shown in red and are represented by warm colors. Colors become progressively cooler for regions with fewer gaze samples.

### Fixation Location

To analyze this in more detail, Figure Figure 3A shows the median X gaze for height, width, and brightness judgments. A repeated measures ANOVA with the within-subject factors of condition (height, width, and bright) and the factor of boosted status (boosted, and un-boosted) reveals a main effect of condition [F(2,28) = 17.73, *p <* .001, 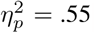, but no main effect of boosted stimuli [F(1,14) = 3.05, *p* = .1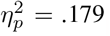, and a significant interaction [F(2,28) = 12.70, *p <* .001, 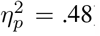. Width and brightness judgments show gaze distributed closer to the center than height judgments (all *p <* .001). However, across all tasks, observers tend to look toward the inner edges of the object closest to each other (all *p <* .05). This pattern holds true for both boosted and un-boosted trials. Notably, for height judgments, when the trial contains an un-boosted object, observers tend to fixate towards the inner edges of both objects to an even greater extent than for boosted trials [*t*(14) = 4.6, *p <* .001].

**Figure 3:**
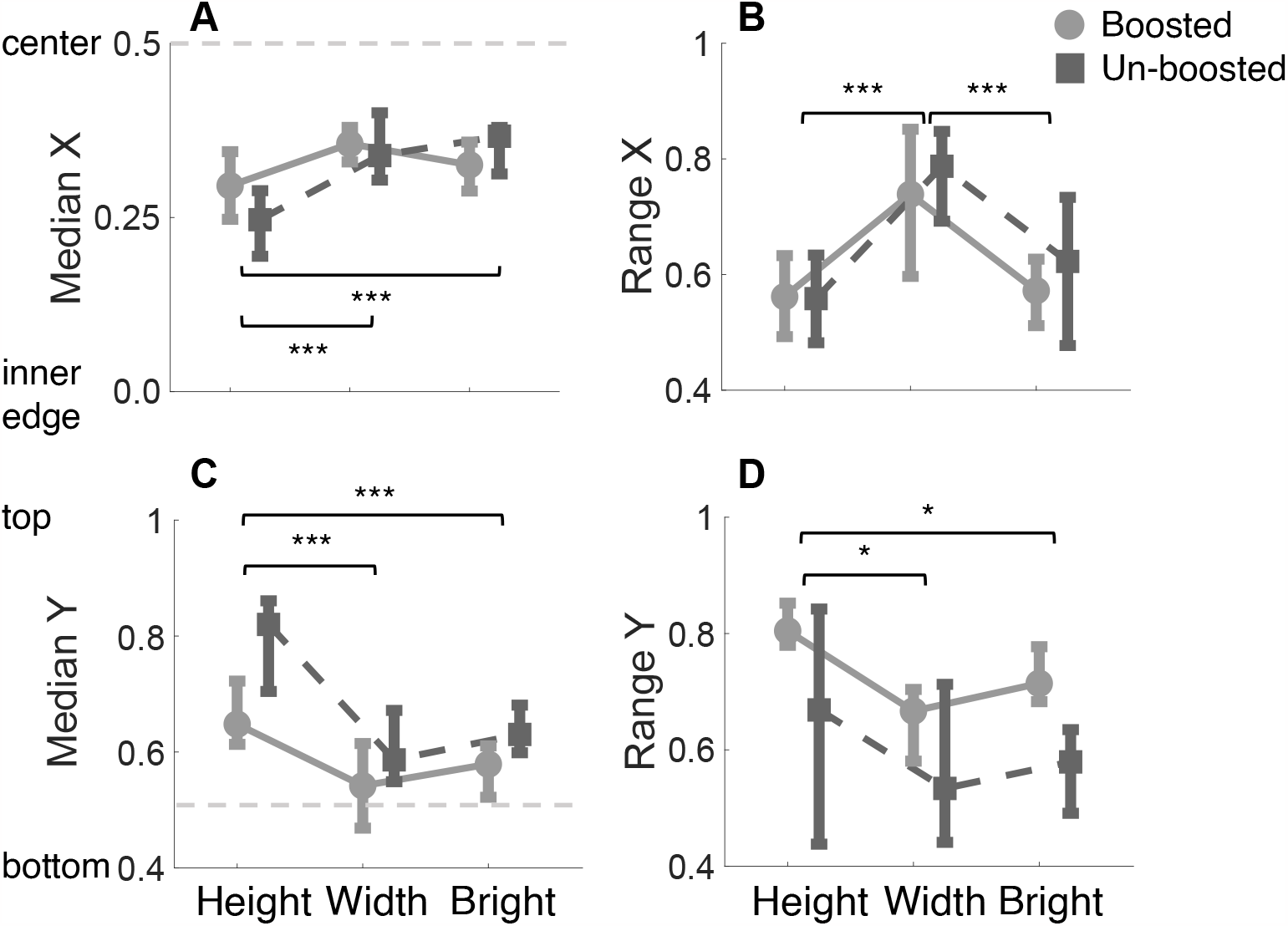
Median gaze and the range of gaze over the objects across tasks. The range quantifies the spread of the distribution of gaze position, which is defined as the difference between the 97.5 and 2.5 percentiles for the X and Y components of gaze. Data are split by whether the trial contained boosted or un-boosted stimuli. The horizontal axis represents the conditions tested. A) and C) present the median X and Y components of gaze, respectively. The dotted grey line represents the object center. B) and D) show the range for the X and Y components of gaze during height, width, and brightness judgments. For all plots, error bars are the interquartile range. If a p-value is less than .05, it is flagged with one star (*). If a p-value is less than .01, it is flagged with 2 stars (**). If a p-value is less than .001, it is flagged with three stars (***).

To quantify the spread of gaze over the object horizontally, Figure 3B shows the range of the X gaze distribution over the objects. The range is calculated as the difference between the 97.5th and 2.5th percentiles for the X and Y components of fixations. A larger value indicates a broader distribution of gaze over the object. We found a main effect of task on the spread of X gaze [*F* (2, 28) = 21.90, *p <* .001, 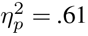, no main effect of boosted stimuli [*F* (1, 14) = 1.52, *p* = .24, 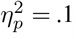 and no interaction [*F* (2, 28) = 1.2, *p* = .32, 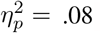. This data reveals a broader spread of gaze horizontally for width judgments as compared to height and brightness judgments. This suggests that for width judgments, observers tend to look towards the left and right edges of the object to a greater extent than for height and brightness judgments.

The vertical component of gaze shows biases depending on the task being performed. To quantify the average vertical component of gaze, Figure 3C shows the median Y gaze for height, width, and brightness judgments. We observe a main effect of condition [F(2,28) = 39.20 *p <* .001, 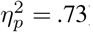, a main effect of boosted stimuli [F(1,14) = 62.01 *p <* .001, 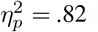, and a significant interaction [F(2,28) = 8.02 *p* = .002, 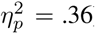. Height judgments show fixations biased toward the top of the object, to a greater extent than width and brightness judgments (all *p <* .001). Across all tasks, fixations in trials with un-boosted stimuli exhibit a vertical bias towards the top of the object. The significant interaction is related to the fact that fixations during height judgment, are in general, directed towards the top of the object to a greater extent than width and height judgments, but this effect is particularly strong for the un-boosted trials.

The vertical spread of gaze is shown in Figure 3D. For condition, we observe a main effect [F(2,28) = 8.44, *p* = .001, 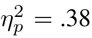, a main effect of boosted stimuli [F(1,14) = 17.06, *p* = .001, 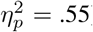, and no interaction [F(2,28) = .554, *p* = .58, 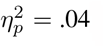. Height judgments show a broader distribution of vertical gaze, as observers are looking towards the top and bottom of the object as compared to width and brightness judgments (*p <* .05). In addition, the introduction of boosted objects led to an overall increase in vertical spread across all tasks.

Width and height judgments require discrimination of surface shape, while brightness judgments comprise a decision about surface properties. Generally, brightness judgments have shown a similar pattern to width judgments for fixation position and distribution. For brightness judgments, are observers fixating on brighter pixels to discriminate the brighter of the two objects? We generated brightness intensity maps for each object shown in the study. Following the analysis pipeline described in the methods section, for each fixation on the object, we calculated the brightness of the pixel fixated. Figure 4 displays the average brightness at fixation for height, width, and brightness judgments. The dotted line represents the average object brightness. Across all tasks, observers tend to fixate on pixels that are brighter than average. The average brightness fixated was significantly different across conditions [*F* (2, 28) = 8.14*p* = .002, 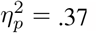, with no main effect of boosted stimuli [*F* (2, 28) = .342*p* = .57, 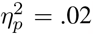, or interaction [*F* (2, 28) = 3.16*p* = .06, 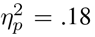. Brightness judgments are associated with fixations on brighter regions of the object compared to width and height judgments (*p <* .01, width vs. height *p* = .5). This finding replicates previous work by (Toscani et al., 2013a), which demonstrated that when asked to judge an object’s lightness, observers tend to fixate the brightest regions on an object, a computationally optimal strategy.

**Figure 4:**
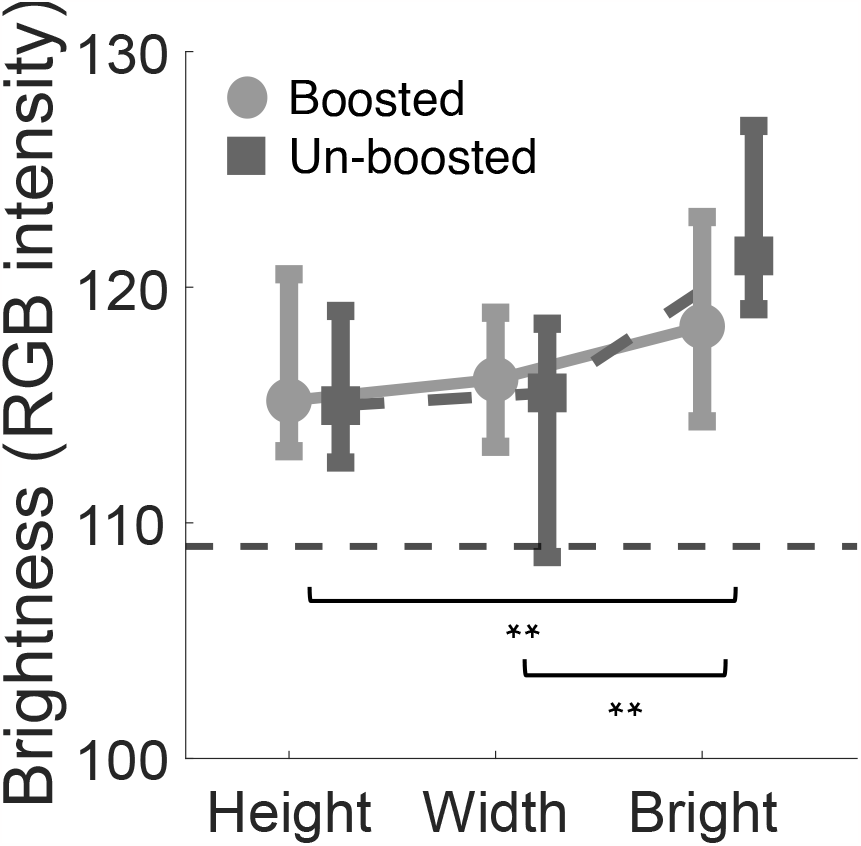
Average brightness at the fixated pixel across different tasks. Brightness is represented as RGB intensity. The dotted line represents the average brightness across all objects tested.

### Fixation Dynamics

We additionally investigated fixation and saccade metrics in order to quantify the temporal dynamics of gaze behavior during the different tasks. The performed task influenced the fixation duration as shown in Figure 5A [*F* (2, 28) = 8.52*p* = .001, 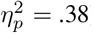, with no main effect of boosted stimuli [*F* (1, 14) = 1.71*p* = .12, 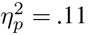, and no interaction [*F* (2, 28) = 2.76*p* = .08, 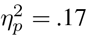. The main effect was explained, by longer fixations during brightness judgments than for width and height judgments (all *p <* .05), but no difference between height and width judgements (*p >* .90). The average saccade amplitude for each task is shown in Figure 5B which does not show a systematic effect of task [*F* (2, 28) = 2.7*p* = .08, 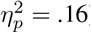, nor a main effect of boosted stimuli [*F* (1, 14) = .56*p* = .46, 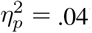, and no interaction [*F* (2, 28) = 0.12*p* = .89, 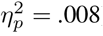].

**Figure 5:**
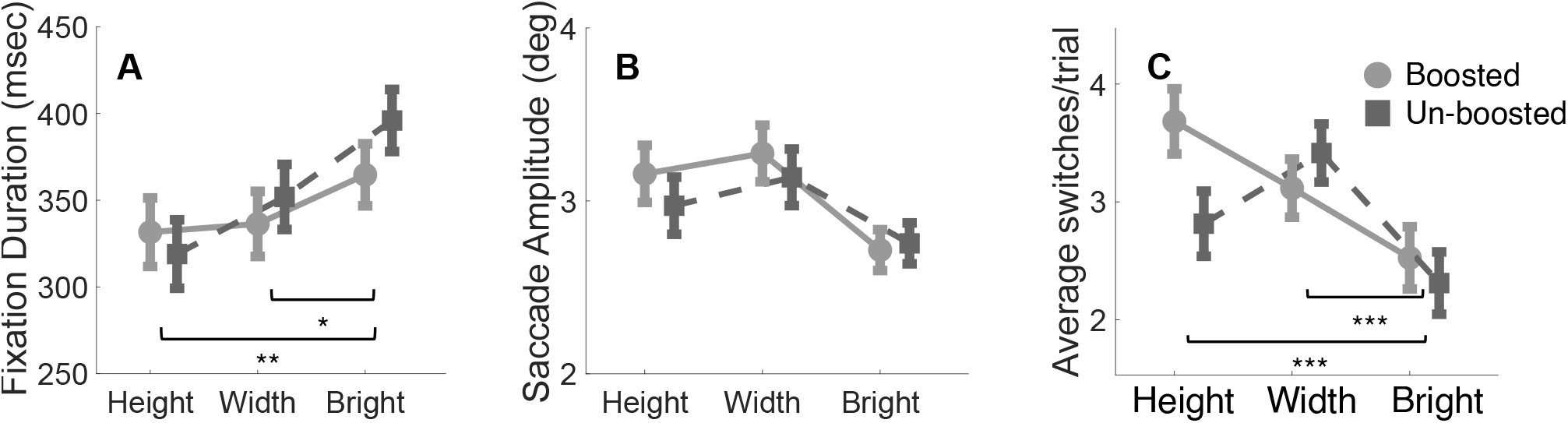
Fixation and saccade metrics for the different tasks. A) shows the mean fixation duration in msec, for each of the tasks. B) Mean saccade amplitude in degrees. C) The average number of switches made per trial between the two objects shown. A switch was classified as a saccade which started on one object and finished on the other object.

As observers are given unlimited time to make these visual judgments, how often they switch between observing one object or another may reveal task-specific biases. Figure 5C shows the average number of switches made during a trial for the different tasks. A switch was considered to take place when a saccade began on one object and ended on the other object. Switch rates show a main effect of task [*F* (2, 28) = 11.54*p <* .001, 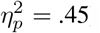], with a main effect of boosted stimuli [*F* (1, 14) = 11.19*p* = .005, 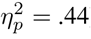], and an interaction [*F* (2, 28) = 8.48*p* = .001, 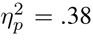]. Brightness judgments are associated with fewer switches than height and width judgments (all *p <* .001). The interaction between task and boosted stimuli for switch rate is primarily driven by height judgments. Height trials that contain boosted stimuli are associated with a higher switch rate than un-boosted trials [*t*(14) = 4.78, *p <* .001]. In the case of un-boosted trials, observers can move their eyes between the closest top corners and directly compare the height of the objects. Such a simple comparison isn’t possible for boosted trials, and for accurate height estimations, observers must repeatedly shift their gaze between both objects to confirm which object is taller. The difference in switch rate between boosted and un-boosted trials did not reach significance for width and brightness judgments (all *p >* .7).

We observed that the type of task influenced fixation dynamics, but are these influences stable across observers? For that we looked at individual differences and how stable these eye movement indices are across visual judgments. For example, if an observer has long fixation durations in one task, do they also have lengthy fixation durations in other tasks? Figure 6 shows the correlations between tasks for fixation duration (Figure 6A-C) and saccade amplitude (6D-F) across tasks. Fixation duration shows a robust correlation between width and brightness judgments (Figure 6A; *r* = 0.65, *p* = .009), height and brightness judgments (Figure 6B; *r* = 0.77, *p <* .001), and height and width judgments (Figure 6C; *r* = 0.67, *p* = .006). Saccade amplitude does not show a positive correlation between tasks (See Figure 6 D-F all *p >* .2). These results suggest fixation duration shows stability across tasks, for width, height, and brightness judgments.

**Figure 6:**
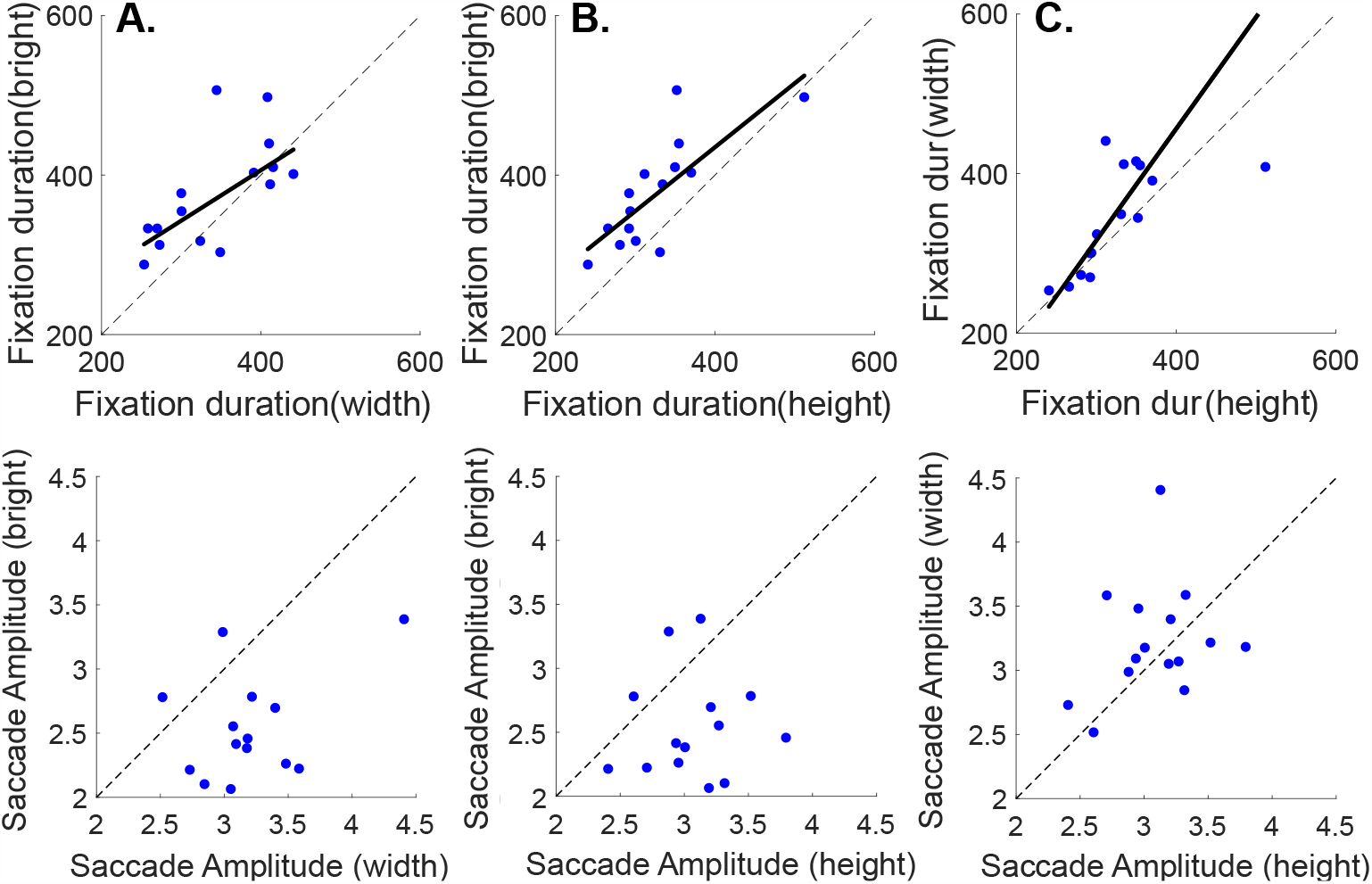
Correlations for fixations metrics between tasks. The upper panel (A,B,and C) shows correlations for the average fixation duration between width, height, and brightness judgments. The lower panel (D,E, and F) show correlations for saccade amplitude (in degrees) between the different tasks. The dotted line represents the 1:1 line; if the data were perfectly correlated, data points would fall on the dotted grey line.

### Summary

The results of the first experiment indicate that there are specific gaze patterns related to performing different perceptual tasks: width judgments lead to fixations with a more central bias and a wider horizontal spread than height and brightness judgments. Conversely, height judgments show fixation biased towards the top of the objects with a wider vertical spread than width and brightness. In the case of un-boosted trials, where both objects were placed on the same surface, observers adopt a strategic gaze pattern. They would shift their gaze between the top closest corners of each object to assess height. This approach minimizes the distance between both objects for comparison. Fixations towards the top of the object are less informative when boosters are used. Conversely, the boosted objects reveal a wider vertical distribution of gaze compared to the trials which contained un-boosted objects. While brightness judgments showed a similar pattern of fixation position and distribution as width judgments, further analysis reveals observers tend to fixate on brighter regions of the object to execute brightness judgments.

Saccade and fixation metrics showed task-specific effects. Brightness judgments were generally associated with longer fixations than width and height judgments. Interestingly, observers also looked back and forth between objects more frequently for height judgments when an object in the scene was boosted. These differences were not seen in the case of width and height judgments. We showed robust correlations within fixation metrics across tasks showing that if an observer makes short fixations during width judgments, they show the same pattern for height and brightness judgments.

Thus, there are task specific adjustments of gaze behavior, but are they also related to performance? In the next experiment, we test a causal link between oculomotor behavior and perception.

## Experiment 2: One object

## Methods Experiment 2

### Participants

Fifteen people participated in the experiment. They were 20-39 years of age (nine identified as female, six identified as male). The experimental protocol was approved by the Institutional Review Board at our university in accordance with the Declaration of Helsinki. Participants signed informed consent forms before participating.

### Apparatus

The VR headset and setup were the same as those used in the previous study.

### Stimuli

Sixty objects were selected from the ArchVizPRO (interior Volumes 6 and 7) asset packs from the Unity Asset store. The standard object used was a 10°×10° white box which remained the same across all conditions tested. During the experiment, observers were asked to judge the width and height of objects that were resized to be 6°, 8°,9°,10°, 11°, 12 °, or 14 ° tall or wide, relative to the standard object. For each height and width size magnitude tested, ten objects were tested, except for the 6° and 14 ° object sizes where five objects were shown.

## Experiment Design

The experiment consisted of two one-hour sessions conducted on separate days. In one session, observers judged the height of objects, while in the other, they judged width. The session order was randomized. The HMD setup and eye tracking calibration and validation procedures were the same as in the previous study. Each session began with observers viewing a standard object. The standard object was always a white box measuring 10° tall and wide (as shown in Figure 7). The standard object was presented on one of three surfaces: a stool, tabletop, or shelf. Observers could view the standard object for as long as they wanted before pressing a key to continue. After this familiarization phase, observers continued to experimental trials, where they were shown a new object. Observers viewed the objects for a duration of two seconds. In the Free Gaze condition, the two-second timer included only those gaze samples that landed on the object, disregarding any gaze samples outside the object. In the gaze-contingent condition, the timer advanced only when the gaze was directed at the fixation region and the object was present. After viewing the object for two seconds, the object disappeared, and the observer responded which object was taller or wider, the standard object or the just-viewed object. To remind observers of the standard object, it was re-shown periodically after several experimental trials. Specifically, at the start of the block of trials, the standard object was shown, and after trials #3, 8, 15, 23, 34, 45. This ensured the standard object was shown at a greater frequency at the start of the block to ensure accurate stimulus encoding. Observers viewed the standard object for as long as they wished before moving on to the next experimental trial.

**Figure 7:**
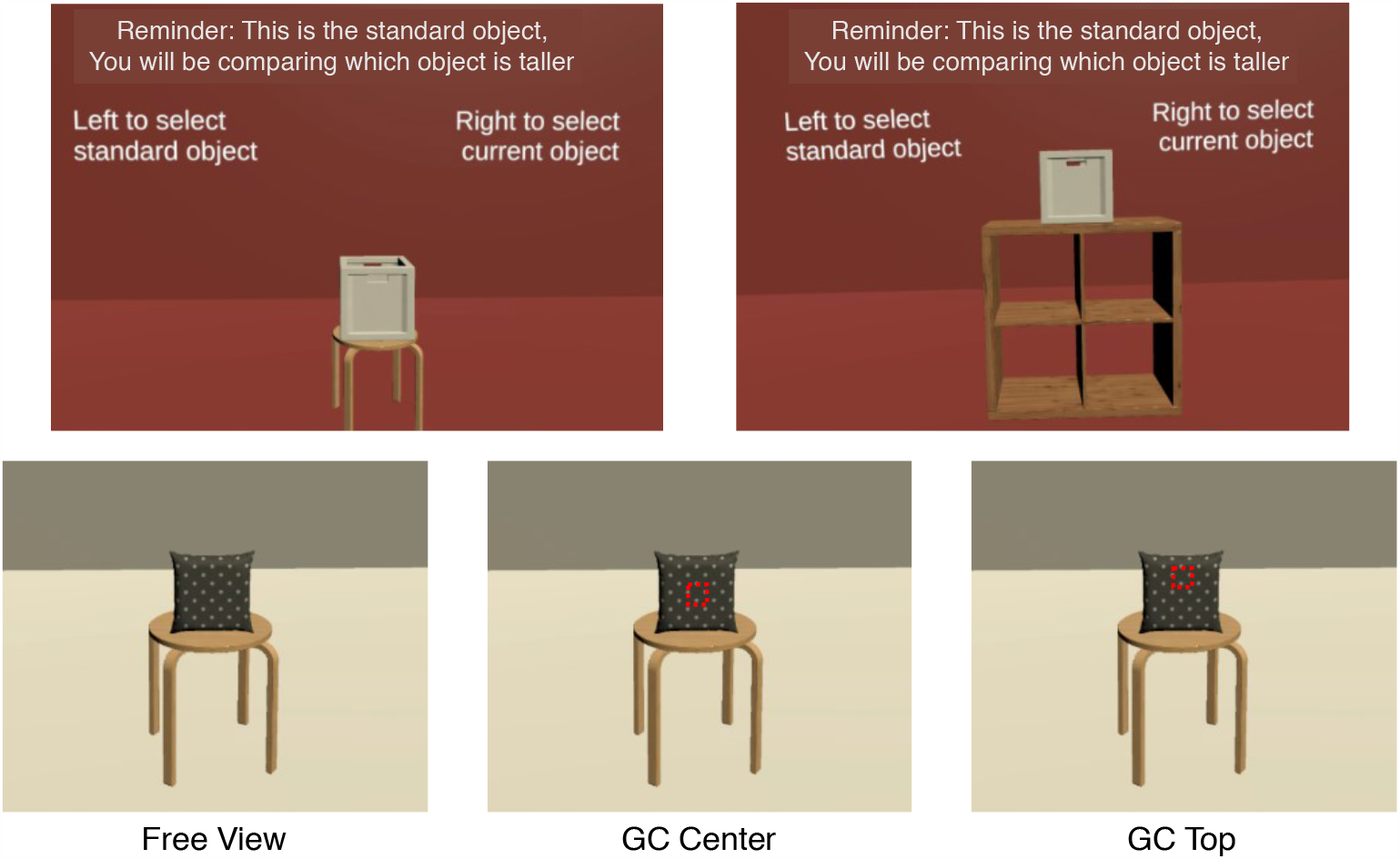
Stimuli and task in Experiment 2. The top panel shows the standard object (a 10°×10° box), which observers could view as long as they wanted at the start of the block of trials. The bottom panel shows an example of the three different viewing conditions tested. In the Free View condition, observers could freely view the object for two seconds. In the Gaze Contingent top and bottom condition, observers were instructed to fixate on the center or top of the object. If their gaze moved outside this fixation region, the object would disappear until their gaze realigned to the appropriate region. The red outlined box represents the fixation window used for illustrative purposes. In the actual experiment, the red cube only appeared to realign gaze after removing the object, in response to fixations falling outside this window.

During each session, observers completed three blocks, a Free Gaze block and two gaze-contingent blocks which forced gaze to specific locations on the object. Each block contained 60 trials, and the same set of virtual objects in the same scenes were used across blocks and tasks, allowing for a comparison of gaze characteristics by task. In the Free Gaze block, observers freely viewed the object shown during the experimental trials for two seconds. If observers looked away from the object, that duration was not counted towards the two-second timer. In the gaze-contingent blocks, each trial started with one of the surfaces (table, shelf, or stool) presented with a 1°x1° red cube above it. Unbeknownst to observers, the red cube was surrounded by an invisible 3°x3° fixation window which remained present during the trial. Observers were instructed to fixate on the red cube. Once gaze remained in the fixation window, the red cube disappeared, and the object for that trial appeared. If gaze stayed within the fixation window, the object remained visible. If gaze moved outside this window, the object would disappear, and the red cube would appear. Observers were instructed to again fixate on the red cube until the object reappeared. The trial ended, and the object disappeared after gaze remained on the object (and in the fixation window) for two seconds in total. Observers then responded which object was taller or wider the object that was just shown or the standard object in memory.

The gaze-contingent setup forces gaze toward specific locations on the object. If where observers fixate determines their performance for height and width judgments, forcing gaze to specific object locations should interrupt performance compared to free viewing. In the gaze-contingent center (Gaze Contingent Center) block, the red cube and fixation window were positioned at the center of the front face of the object. In contrast, for the gaze-contingent top (Gaze Contingent Top) block, the fixation window, and the cube was positioned at 70% of the object’s height. These positions were used to approximate the average gaze positions for the height and width task from the previous study. The horizontal and vertical positions of the fixation window and cube were randomly jittered by +/0.2, 0.4, 0.6, 0.8, or 1°. Block order on each day was counterbalanced using a Latin Square Design.

In this experimental design, the free-viewing task measures natural behavior, allowing observers to execute their preferred gaze strategy. The Gaze Contingent Center block forces fixation towards the object center, interrupting the typical strategy of looking at the top of the object during height judgments. This may lead to poorer performance in the height task as compared to the width task, where observers naturally fixate centrally on the object. Enforcing fixation centrally may not have as much impact on performance for width judgments. Similarly, the Gaze Contingent Top block may lead to poorer performance in the width task while potentially preserving performance on the height task, as fixation is directed towards the top of the object, where observers are likely to naturally fixate.

## Data Analysis

Data analysis procedures were similar to those described in the previous experiment. Raw gaze positions were saved during each trial for XYZ coordinates and were further processed offline. Object information, such as the collider label and location details, was also saved. For the Free Gaze condition, gaze samples that were part of a fixation and fell on an object were normalized between 0-1 based on the object size. In order to quantify discrimination performance between the Free Gaze and Gaze Contingent conditions, psychometric functions were fit to the data. Psychometric functions were estimated by calculating the proportion of ‘standard wider’ or ‘standard taller’ responses for each stimulus size. Stimulus size was represented as the percent difference in width or height from the standard object. We calculated separate psychometric functions for each block of the experiment (Free Gaze, Gaze Contingent Top, and Gaze Contingent Center) for both width and height judgments, using the Psignifit toolbox (version 4 for Matlab (Schütt, Harmeling, Macke, & Wichmann, 2016)). We used a cumulative Gaussian distribution with four parameters, including the point of subjective equality (PSE), the sigma of the function, guess rate, and lapse rate. Discrimination performance was quantified by the σ parameter of the cumulative gaussian function. All statistical analyses were performed in JASP.

One observer was excluded from the analysis as their sigma values were greater than the mean + 2 standard deviations of all observers, for the Free Gaze and Gaze Contingent Top conditions.

## Results Experiment 2

The goal of experiment 2 was to investigate the causal link of task-specific oculomotor behavior on perceptual judgments. Specifically, we will contrast performance on width and height discrimination tasks when observers freely viewed the object (executing their preferred gaze strategy) to gaze-contingent conditions, where participants are forced to fixate on a specific location on the object.

To confirm our gaze contingent manipulation was executed effectively, Figure 8 shows the average gaze heat maps for both height and width judgments, split by free viewing and gaze contingent conditions. Figure 8A and D show that gaze is restricted towards the top of the object for height judgments, and that observers tend to look towards the object’s center for width judgments. If the gaze-contingent manipulation effectively restricts gaze, gaze should be constrained towards the center for Gaze Contingent Center trials, and towards the top for Gaze Contingent Top trials. Figure 8B and E show data from the Gaze Contingent Center condition for height and width judgments, while Figure 8 C and F show data from the Gaze Contingent Top condition for height and width judgments. The Gaze Contingent Center heatmaps show gaze restricted centrally, while the Gaze Contingent Top heatmaps show gaze shifted towards the top of the object. Importantly, the spread of the data is restricted, especially compared to the Free Gaze conditions, suggesting the gaze-contingent manipulation was successful.

**Figure 8:**
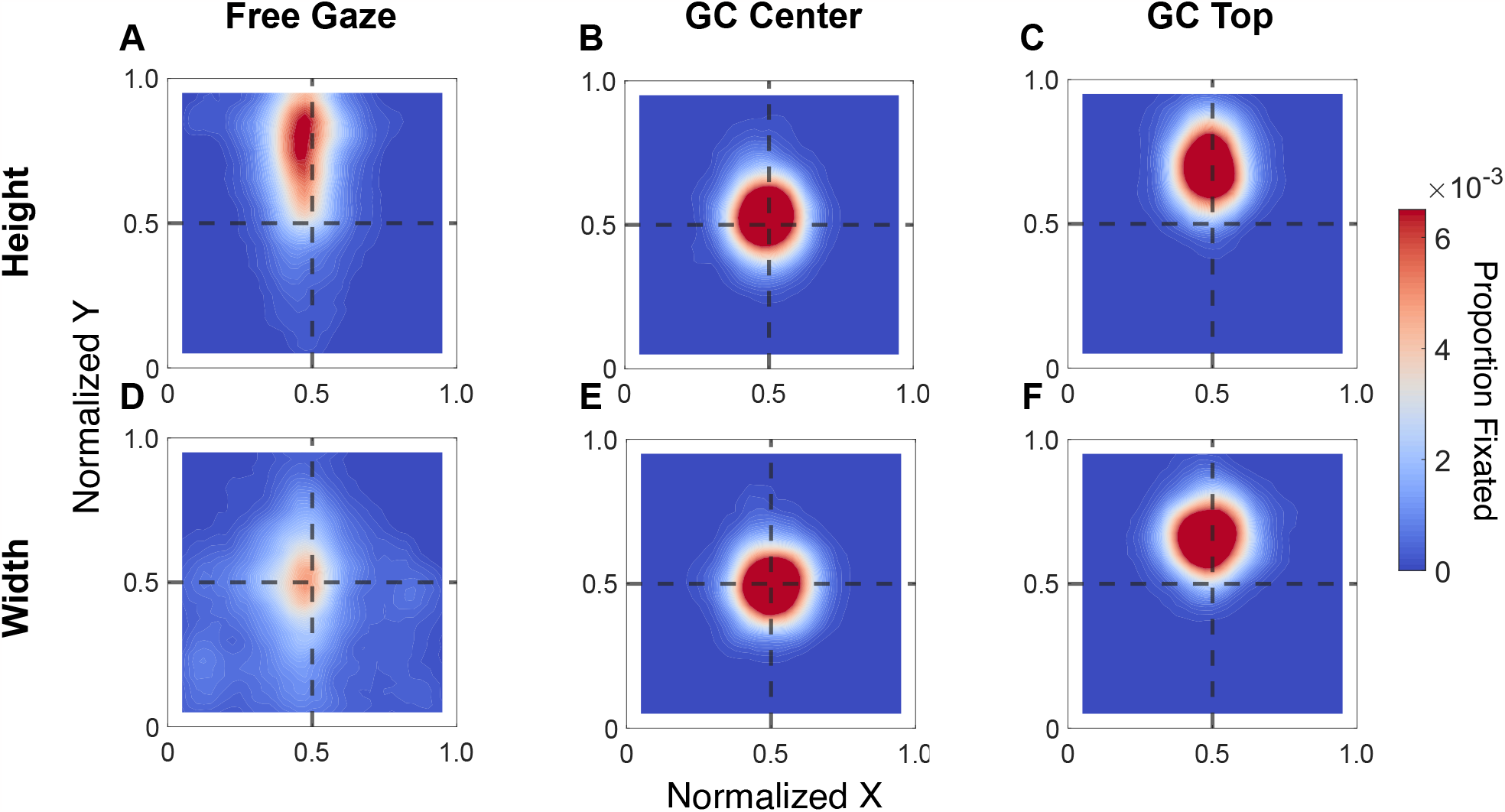
Average heat map for height judgments (top row, A-C) and width judgments bottom row D-F) judgments. Gaze data are split between free viewing and gaze-contingent conditions. The heatmap is relative to object position; values that fall close to the top represent fixations biased toward the top of the object, while values close to the center represent fixations biased centrally.

### Visual discrimination performance

To assess discrimination performance, psychometric functions were fit to the data. Figure 9 shows the psychometric fits for one representative observer in the Free Gaze, Gaze Contingent Center, and Gaze Contingent Top conditions for width judgments. Notably, the psychometric functions of this particular observer do not show systematic differences between the free viewing and gaze-contingent conditions for either width or height judgments; the discrimination threshold is constant between these three psychometric functions.

**Figure 9:**
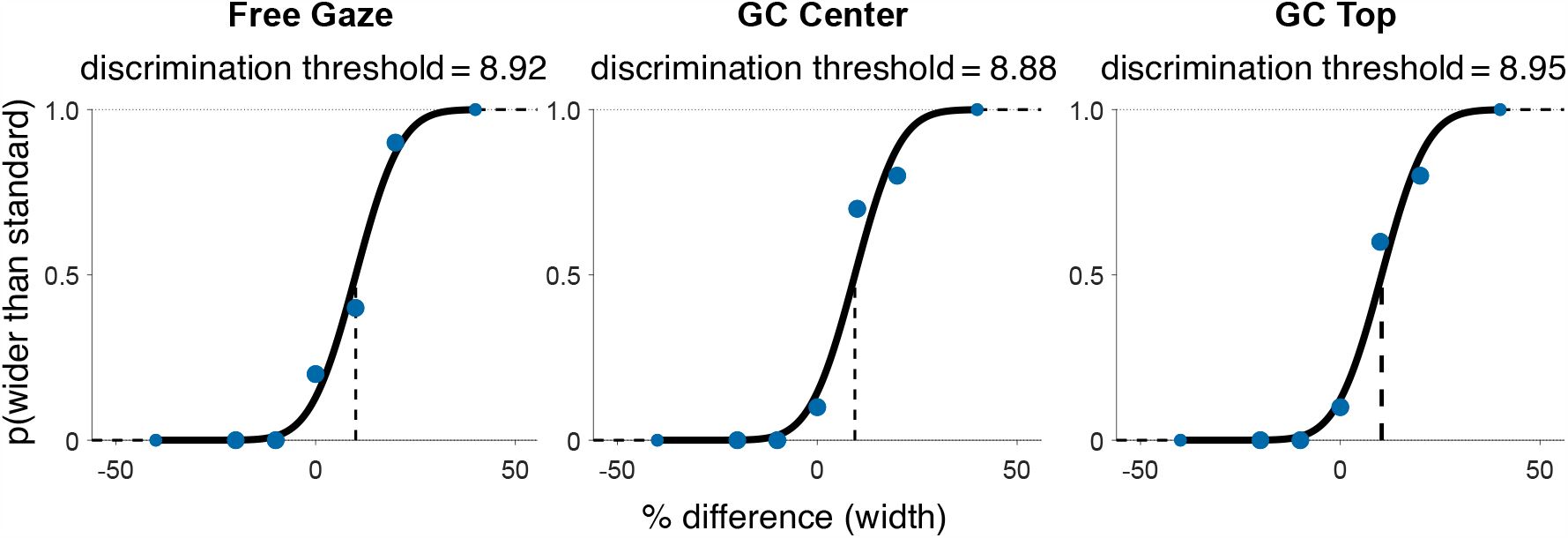
Psychometric fits for data from a single representative observer. Data are fit with a cumulative Gaussian function, and a vertical line marks the 50% point of subjective equality (PSE). Three separate psychometric functions were fit for data from each of the conditions tested. The left panel shows data from the Free Gaze condition, the center panel shows data from the Gaze Contingent Center condition, and the plot on the right shows data from the Gaze Contingent Top condition.

Figure 10 presents the average discrimination threshold for width and height judgments across all observers. The data are split by viewing condition: Free Gaze, Gaze Contingent Center, and Gaze Contingent Top. A one-way repeated measures ANOVA reveals no significant difference in discrimination performance between these groups for width judgments [F(2,28) = 0.35 *p* = .71, 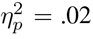] and height judgments [F(2,28) = 0.175, *p* = .840, 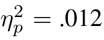]. This means discrimination performance is consistent across the viewing conditions. The gaze-contingent manipulation, which was designed to interrupt the preferred gaze strategy used by observers, did not disrupt discrimination performance for width and height judgments.

**Figure 10:**
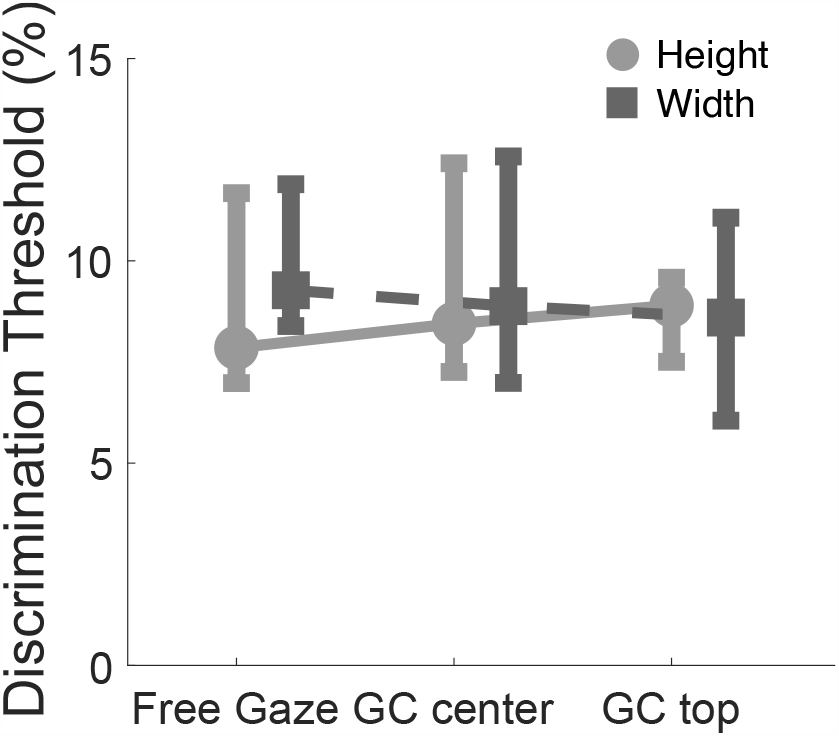
Effect of viewing conditions on discrimination performance. Median discrimination threshold by the gaze conditions tested for width (dark grey) and height (light grey) judgments. Data are split between the free gaze, Gaze Contingent center, and Gaze Contingent top conditions. Smaller discrimination thresholds represents better discrimination performance.

### The potential role of gaze dynamics

One reason for the lack of impairment due to the gaze contingent condition could be that while in the gaze continent condition observers fixated at a specific location, in the free gaze condition they could actively move their eyes. We know that during eye movements visual sensitivity is impaired (Binda & Morrone, 2018), so the active movement might also harm discrimination performance. To investigate this effect, we computed correlations between gaze metrics (saccade amplitude and fixation duration) with discrimination threshold for height and width judgments for the free-gaze condition (see Figure 11). If oculomotor dynamics indeed influence discrimination performance, there should be a correlation between fixation/saccade dynamics and discrimination performance. Our results don’t show a significant correlation between saccade amplitude and discrimination performance, or fixation duration and discrimination performance (all *p >* .49, all *r <* .24). This seems to suggest, that the lack of a difference in performance for the gaze-contingent condition is not related to negative impacts of gaze dynamics in the free-viewing condition.

**Figure 11:**
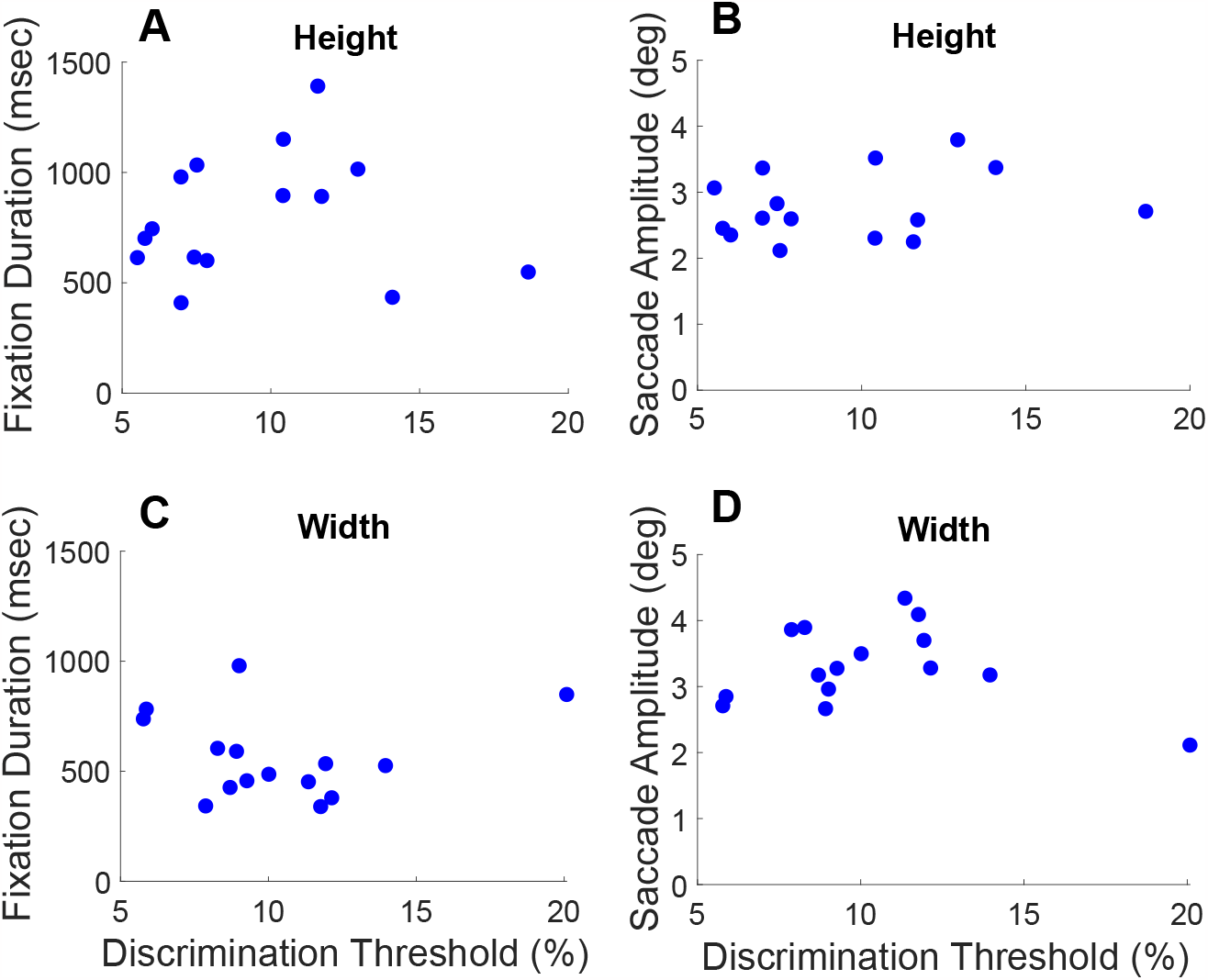
Correlations for gaze metrics and discrimination threshold. A) and C) show the correlations between fixation duration and discrimination threshold for height and width judgments. B) and D) show the correlation between saccade amplitude and discrimination threshold for height and width judgments.

### The potential role of object size

Another reason why our gaze contingent manipulation could have no effect, is that the size of some objects might have allowed observers to process the whole object during one central fixation. To look into this possibility, we analyzed the fixation positions across the different object sizes. If discriminating size for larger objects necessitates distinct fixation strategies as compared to smaller objects, we would observe these differences in the Free Viewing trials. Figure 12 plots the average relative XY gaze position by object size. Note that the XY coordinates correspond to object proportions, where 0,0 represents gaze towards the bottom left of the object and 1,1 towards the top right. As in Experiment 1, observers show a consistent pattern of gaze; when discriminating width, fixation is biased towards the center of the object, while when judging height, fixations are directed at 70% of the objects’ height. This strategy is remarkably consistent across different object sizes and remains robust whether the object is 6° or 15° in size. Observers appear to be dynamically adjusting their fixation location to a fixed proportion of the object’s size. Notably, this means two things: (1) observers do not adjust their behavior based on object size, since they prefer to look at a fixed proportion of the object and not, for example, at a fixed distance relative to the edges or center of the object, and (2) observers must automatically acquire an accurate representation of object size to achieve fixation on a fixed proportion of the object so consistently.

**Figure 12:**
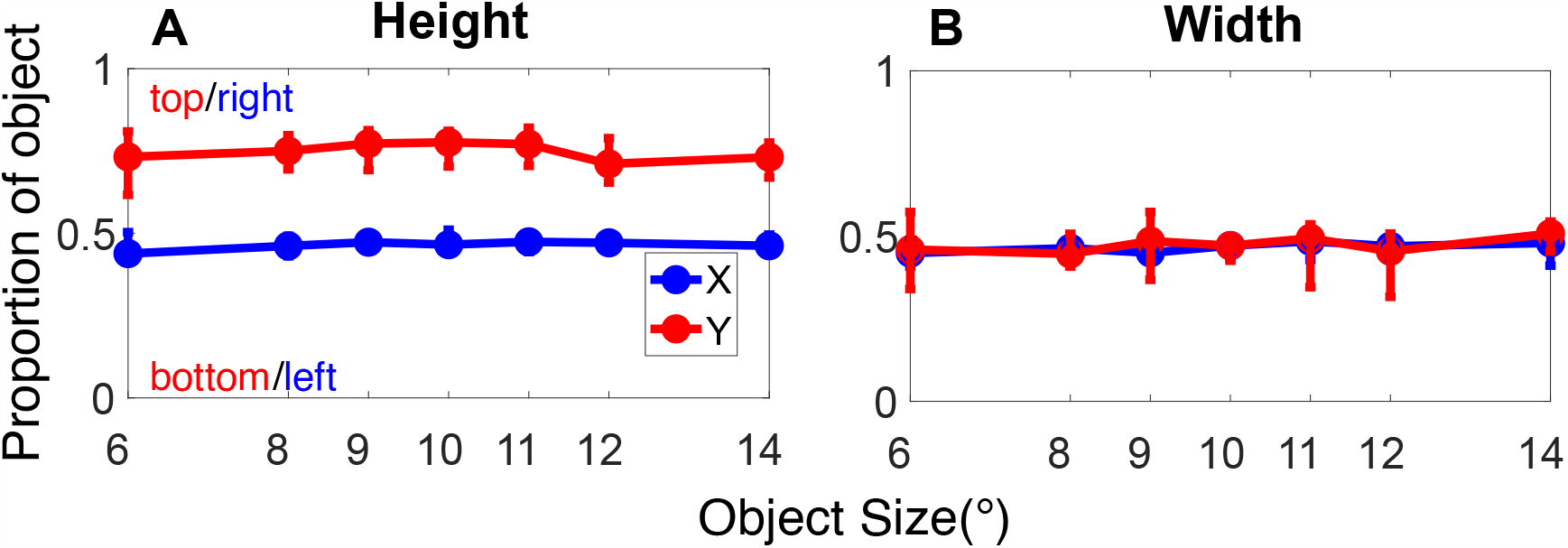
Average gaze location relative to the object. Normalized XY gaze position, where the coordinates correspond to the object proportions. 0,0 represents gaze falling toward the bottom left of the object, while 1,1 represents gaze toward the top right of the object. Data for width judgments are shown in the left panel, and height judgments are shown in the right panel.

### Fixation Dynamics

To characterize patterns of oculomotor behavior, Figures 13A shows the average fixation duration, and 13B shows the average saccade amplitude made for height and width judgments. When making height judgments for single objects in comparison to a memorized standard observers tend to make longer fixations [*t*(14) = 2.7, *p* = .02] and smaller saccades [*t*(14) = −2.5, *p* = .02] as compared to width judgments. This is in contrast to the results of experiment 1, where two objects were directly compared, which overall led to much shorter fixation durations (compare Figure 13 and Figure 5).

**Figure 13:**
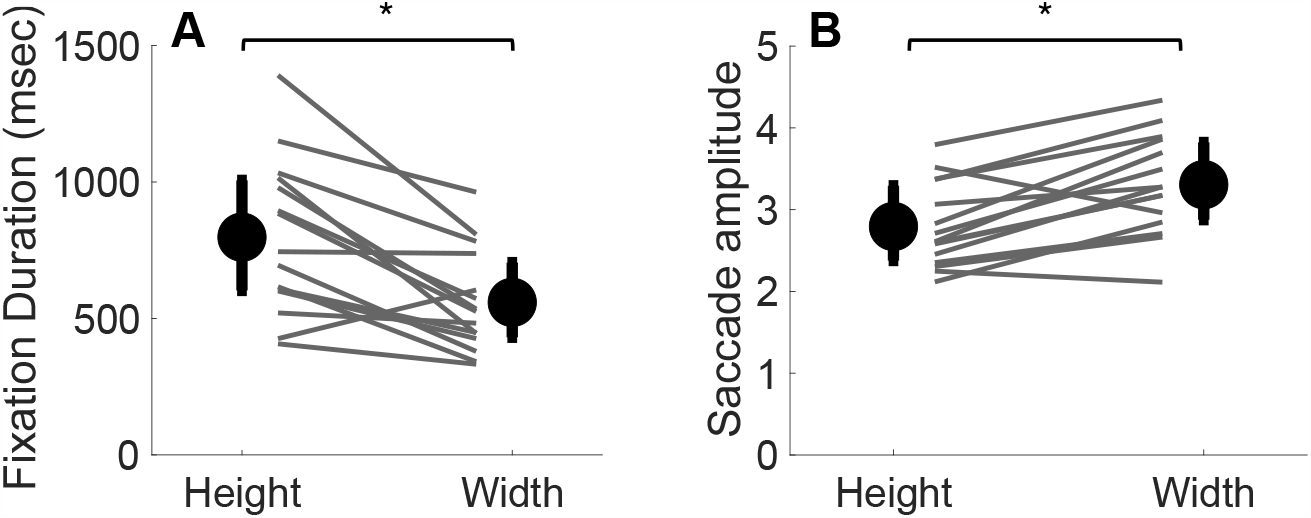
Fixation and saccade dynamics for height and width judgments. A) shows the average fixation duration in msec, and B) shows the average saccade amplitude.

As in the previous experiment, we also looked at the consistency of behavior across tasks by correlating individual differences in fixations frequency and saccade amplitude across tasks. A positive correlation between width and height judgments for fixation duration means participants with long fixations in one task tend to have longer fixations in the other task as well. The analysis in Figure 14A shows the correlation between the fixation duration made during height and width judgments and Figure 14B shows the correlation between saccade amplitude for both tasks. Fixation duration for height and width judgments shows a robust positive correlation [*r*(13) = .68, *p* = .006]. The amplitude of saccades made during the height and width task also shows a strong and positive correlation [*r*(13) = .71, *p* = .003]. This suggests individual differences in fixation duration and frequency are consistent across surface shape judgments.

**Figure 14:**
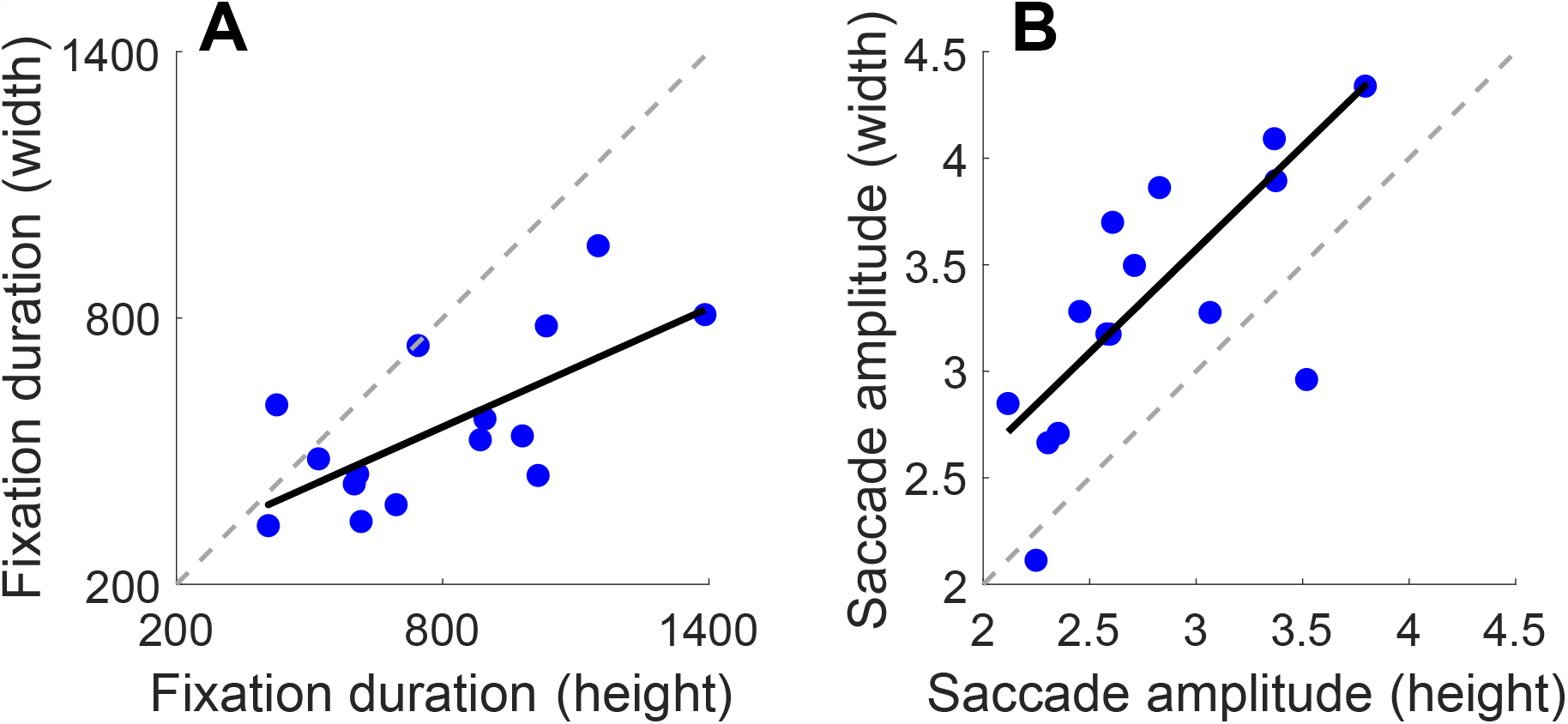
Correlations for fixations metrics between tasks. A) shows correlations for the average fixation duration between width and height judgments. B) show correlations for saccade amplitude between the different tasks. The dotted line represents the 1:1 line; if the data were perfectly correlated, data points would fall on the dotted grey line.

### Summary

We tested a causal link between oculomotor behavior and perception in the current experiment. Observers freely viewed an object or viewed it with a gaze-contingent set up to force fixation to a specific location. In the free gaze condition, width judgments were accompanied by fixations biased towards the object center. Conversely, height judgments showed gaze shifted towards the top of the object, replicating experiment 1, which presented two objects. Surprisingly, discrimination performance showed no difference between free gaze and gaze-contingent conditions for width and height judgments. This effect could not be explained by potential negative influences of gaze dynamics nor a change in behavior for different object sizes.

## Discussion

We have demonstrated that gaze behavior dynamically adapts to a given perceptual task and the presented visual stimuli. However, despite these robust, and adaptive oculomotor strategies, we found no causal influence on eye movement behavior for task performance. In the following sections we will discuss the implications of these results for why observers adjust their gaze, and how they are related to perceptual performance.

### How does gaze adapt by task?

When observers are presented with two objects and asked to judge which is taller, wider, or brighter, they look at different parts of the object, adapting their gaze to the task at hand. We initially hypothesized that observers could use their gaze as some kind of measuring tape to provide additional cues for the following perceptual judgments, however we observed that they rather seemed to fixate at different locations on the object: When comparing the width of two objects, fixations are biased towards the center of the object and shows a greater horizontal spread. Moreover, when judging height, observers tend to shift their gaze toward the top of the object, and a broader vertical spread. Interestingly, when given the option to directly compare the height of two objects sitting next to each other on the same surface, observers look back and forth between the two objects, comparing the top closest corners. This is an adaptive and optimized strategy, as energy expenditure is minimized, allowing a direct comparison between the objects. This ‘switching’ behavior changed and was more pronounced when the surfaces the objects were placed on were uneven, preventing direct comparison of the top corners of the object. In this case, observers alternated their gaze between both objects often, likely to better compare the absolute height of the objects. The boosted condition mainly affected height judgments, leading to more uncertainty, and an increased number of switches. This change in switching suggests that under higher uncertainty and the lack of an easy comparison, observers tend to sample both objects more frequently to ensure their perceptual decision. Interestingly, when comparing the results of experiments 1 and 2, we observed that when only one object was visible and needed to be compared to a memorized standard, the overall typical fixation locations stayed the same, but fixation duration become longer. Thus, even here, when observers didn’t need to sample two objects, but only needed to gather information about one, they decided against the use of an systematic scanning of the object to measure their size. In contrast, observers rather spent the time by increasing their fixation duration on the preferred locations to gather more detailed information about the current object of interest. For brightness judgments, observers tend to fixate on brighter regions of the object, a finding which replicates prior work (Toscani et al., 2013a), and represents a computationally optimal strategy. These results show that the given task influences eye movements and fixations patterns, a finding which reproduces previous work (Brouwer et al., 2009; Chen et al., 2021; Rothkopf et al., 2007; Tatler, Hayhoe, Land, & Ballard, 2011; Yarbus, 1967).

### How could eye movements influence size judgments?

When considering the usefulness and potential role of eye movements in improving visual judgments, the oculomotor system could provide valuable additional information. Making an eye movement from one side of the object to the other could provide an additional cue to its size. Information about the length of an eye movement is accessible from two different sources: First, mechanoreceptors in the muscles sense the position of the eyes (Sun & Goldberg, 2016; Sherrington, 1899, 1918) and provide proprioceptive feedback (Donaldson, 2000). Second, information may also arise from the Efference copy generated from the innervations to the extraocular muscles. The Efference copy is an internal copy of a motor innervation, and acts as an extraretinal source of information for perceptual localization and motor activity (Bridgeman, 1995). Both sources of feedback from the oculomotor system are used and combined to estimate the current eye position (Poletti, Burr, & Rucci, 2013), and this could be utilized to estimate measures of object size.

As opposed to an oculomotor strategy which would utilize proprioceptive and Efference copy information, our results showed observers typically tend to fixate at specific locations on the object. Selecting an optimal location on the object could lead to better processing of sensory signals, as the fovea is aligned at the object center. This may have inadvertently improved performance in the gaze contingent case. Previous work eye tracking elite athletes has documented ‘Quiet Eye’ behavior. This arises as athletes tend to fixate on regions of interest earlier and for longer durations than novices (Vickers, 2007), which is then related to better perceptual judgements and interception behavior. Observers seem to rely on a similar approach in our perceptual tasks, since they rather choose to fixate for a longer time on a specific location than actively sampling the whole object. This could also explain, why the gaze contingent manipulation did not impair performance, since it biased oculomotor behavior towards longer fixations.

In line with the ‘Quiet Eye’ idea, making a lot of large eye movements could have also impaired perceptual performance. We know that visual sensitivity is impaired around the time of saccadic eye movements (Binda & Morrone, 2018) and that during critical moments in time (e.g. when intercepting a moving target), observers sometimes even actively suppress the execution of a saccadic eye movement (Goettker, Brenner, Gegenfurtner, & Malla, 2019). This could also explain the lack of difference between the free-gaze and gaze contingent condition, due to an impaired performance in the free-gaze condition. However, this explanation is unlikely: One would expect larger saccades in the free gaze condition to be associated with worse performance if this interpretation was correct, and we do not find such a relationship between discrimination performance and saccade amplitude (as shown in Figure 11).

In summary, although active sampling behavior could provide essential additional information for judging object size, our results suggest that basic height and width judgments primarily rely on perceptual coding rather than an oculomotor strategy.

### Relationship between fixation routines and visual judgments

The current experiments show that although fixation behavior shows robust, task-specific biases, interrupting these routines doesn’t affect discrimination performance. This highlights the role of perception in estimating surface shape measurements and reveals the visual system is not dependent on a fixed oculomotor strategy to perform such evaluations. One strength of the perceptual strategy is that it is more robust: If the oculomotor system were measuring the size of the object using gaze (a measuring tape approach), the resulting size estimation would be in visual degrees. However, visual angle alone is inadequate to estimate size correctly, as many combinations of size and distance can produce equivalent retinal size (Sousa, Brenner, & Smeets, 2011; Sousa, Smeets, & Brenner, 2012). Although familiarity and familiar size may provide a strong cue to object size (McIntosh & Lashley, 2008; Gillam, 1995), the retinal angle alone is likely an insufficient cue for object size judgments.

Another factor related to the size of the objects, is that the tested objects were relatively large; observers did not necessarily need to foveate the object to acquire an impression of the object’s size. This is not true of all tasks; if fine spatial detail is required for discrimination, gaze has a causal role, and the stimulus must align on the foveola for the task to be performed (Ko, Poletti, & Rucci, 2010). In this case, there is a causal and critical role of gaze; fine eye movements help optimize performance by shifting the eyes so the stimulus falls on the fovea (Clark, Intoy, Rucci, & Poletti, 2022; Intoy & Rucci, 2020). This is a unique feature of vision and suggests eye movements are uniquely adapted to the given task, and are recruited when necessary. Gaze may not have been relevant for our task, as observers can encode an impression of object size without precisely aligning the fovea on the object. Such nuances in the influence of task between the visual and motor systems lead to complexities in directly comparing the two.

If the fixation biases we found represent an optimal strategy, why did the gaze-contingent condition not affect the discrimination threshold? One possibility is that scene gist processing contributes to the task. Previous work has shown that very brief exposures of scenes (50-300 msec) are sufficient to understand and process the gist of an image and to categorize it at a basic level (Bacon-Macé, Macé, Fabre-Thorpe, & Thorpe, 2005; Fei-Fei, Iyer, Koch, & Perona, 2007; Gillam, 1995; Larson & Loschky, 2009). Additionally, Larson et al., (2009) found that peripheral vision plays a crucial role in gist perception, and is more useful than high-resolution foveal vision at recognizing the overall meaning of the scene. In our task, it is plausible that a coarse gist-like representation of the object is utilized for size judgments. This could enable a fast representation of object size, irrespective of where observers are fixating on the object. From the current studies, basic visual size estimations appear to rely on a perceptual strategy instead of a motor strategy. In contrast, Lederman et al., (1967)’s work reveals haptic discrimination relies on clear motor routines, which, when interrupted, reduce performance. However, the haptic system is markedly different from the visual system. Object encoding is largely external, as observers must touch the object, while visual encoding is primarily an internal process. Some features of objects can only be accessed through a haptic interaction. For example, the weight of an object can’t be properly assessed till the object is lifted, or there are fine-scale textures that are imperceptible to sight, but which can easily be felt. In addition, a single visual fixation allows the processing of multiple objects. In contrast, a comparable brief haptic exposure is typically restricted to a single object, instead requiring multiple movements to integrate information about the object (Metzger, Lezkan, & Drewing, 2018; Lezkan, Metzger, & Drewing, 2018). In line with the gist idea, our results show that across object sizes, observers consistently gaze at a fixed proportion of the object, which indicates observers rapidly acquire an accurate representation of object size. Such a fine representation in the haptic domain requires longer exploration of the object and necessitates exploratory movements specific to the task queried.

Although gaze strategy has no causal influence on visual judgments for width and height, this finding may be specific to low-level size judgments. A distinction can be made between low-level features, such as size, and more complex material properties, such as lightness. Object size is considered to be a low-level visual feature, processed in the primary visual cortex (cortical area V1). The retinotopic organization of V1 lends well to the encoding of spatial properties, and previous work has shown evidence for V1’s role in object size perception (Schwarzkopf, Song, & Rees, 2011) and the integration of retinal size and depth cues (Fang, Boyaci, Kersten, & Murray, 2008). In contrast, judging an object’s lightness requires a complex integration across the object. Observers must pick a single ‘average’ intensity value for the whole object, across a distribution of non-uniform intensity levels. Current theories suggest lightness perception is a mid-level visual feature requiring multiple physiological mechanisms at different stages of the visual hierarchy (Gilchrist, 2007; Kingdom, 2003; Boyaci, Fang, Murray, & Kersten, 2007; Rossi, Rittenhouse, & Paradiso, 1996). As visual acuity, contrast sensitivity, and color sensitivity vary with retinal eccentricity (Virsu & Rovamo, 1979; Weale, 1953), the visual system must further integrate this representation from multiple samples. Our results show that when asked to compare which of two objects is brighter, observers don’t randomly choose regions within the object, but instead fixate brighter regions to make their judgments. This replicates previous findings by Toscani, Valsecchi & Gegenfurtner (2013), and represents a computationally optimal strategy. Their work additionally showed that when using a gaze-contingent paradigm to force observers to fixate on either a dark or light region of the object, their brightness matches were biased towards the brightness of the region fixated. The brightness of the region fixated directly influences brightness judgments. This suggests there is an optimal way for observers to sample the object with their fixation strategy. Our work additionally found longer fixation durations for brightness judgments than for height and width judgments. Local brightness varies over the object and must be investigated in great detail. In such a detail-oriented task, gaze allocation matters, and observers are fixating on specific, task-relevant locations for longer. In contrast, low-level features such as height and width judgments can be solved perceptually; fixation location doesn’t influence discrimination performance. In this case, it may be useful to fixate at any location on the object to get a coarse representation of shape.

Future directions include understanding the role of fixations in perceptual judgments for complex material properties. When object features directly contribute to visual judgments, oculomotor behavior may reveal specific patterns to exploit these regularities. In the future, modulating task complexity may reveal further oculomotor strategies. Although the current environments tested contained photorealistic objects, more clutter is typical of real-world environments. Additionally, the tasks used in the current study were relatively simple, involving one or two objects on a surface, in an otherwise empty room. The simplicity of the task may have limited the effectiveness of the gaze-contingent manipulation, as observers did not need to execute complex oculomotor routines to complete the task. It is essential to consider that the human visual system is active by nature (Ballard, 2021; Findlay & Gilchrist, 2003; Noë & O’Regan, 2001). To gain deeper insights, we plan to design tasks that involve more dynamic and complex environments, integrating the visual system in a way that better reveals exploratory routines critical for each task. This approach will likely provide more valuable information about how the visual system functions in diverse and challenging scenarios.

### Fixation metrics show stability across tasks

We observed consistent individual differences for fixation duration and saccade amplitude across tasks, which replicates previous findings (Castelhano & Henderson, 2008; Rayner, Li, Williams, Cave, & Well, 2007; Andrews & Coppola, 1999; Poynter, Barber, Inman, & Wiggins, 2013; Zangrossi, Cona, Celli, Zorzi, & Corbetta, 2021). Specifically, fixation duration shows intra-observer stability across tasks: observers with long fixations in one task will have long fixations in another. Previous work has additionally shown fixation duration in scene perception, and face-processing tasks are highly correlated (Castelhano & Henderson, 2008; Rayner et al., 2007). This work also showed high correlations between fixation duration when observers view line drawings, color photographs, and color renderings of 3d models of scenes. Previous work has additionally shown correlations between saccade amplitude for reading and visual search tasks (Andrews & Coppola, 1999). These previous studies show a great deal of consistency within individuals when comparing across different passive viewing tasks on a computer screen. Our experiment extends these results now to photorealistic VR. In addition, unlike these previous studies, which tested different tasks and stimuli, our stimuli were the same across tasks, meaning low-level stimuli characteristics were equivalent across tasks. This again points to the importance of looking beyond averages and at individual behavior: Although eye movements differ between individuals, these differences are more than measurement noise.

## Conclusions

The current experiments investigate the gaze strategy executed for different perceptual tasks. The experiments were presented in photorealistic VR environments, with naturalistic objects. Observers made basic visual judgments on object shape and surface properties while their gaze was tracked. Tasks were performed with the same set of virtual objects in the same scene, allowing for a comparison of gaze characteristics for each task. The results show that although gaze behavior shows adaptive behavior when observers freely view the objects, when these chosen gaze behaviors are interrupted with a gaze-contingent paradigm, discrimination performance remains largely unchanged. These fixation biases, although robust between observers and within a task, do not appear to exert a causal influence on basic visual judgments. This demonstrates that visual system seems to rely on a perceptual strategy in estimating surface shape measurements and reveals that the visual system does not have to rely on an oculomotor strategy to perform such evaluations.

## Notes

### Competing Interest Statement

The authors have declared no competing interest.

